# Recovery of structural integrity of epithelial monolayer in response to massive apoptosis-induced defects

**DOI:** 10.1101/2022.08.08.503238

**Authors:** Yuan Yuan, Yu Long Han, Jing Xia, Hui Li, Chao Tang, Fangfu Ye, David A. Weitz, Ming Guo

## Abstract

Apoptosis exists ubiquitously in organisms and plays an essential role in maintaining the homeostasis of functional tissues. While the signaling pathway of cell apoptosis has been widely studied, the mechanism of how apoptotic cells regulate the structural homeostasis of a living tissue still remains largely elusive. Using a functional epithelial monolayer as a model system, we find that the integrity of the epithelium is interrupted by apoptosis-induced defects, with an increasing permeability to small molecules across the epithelium. The defects promote a structural reorganization through enhanced cell spreading and migratory dynamics, resulting in a quick recovery of epithelium integrity. Moreover, we show the epithelial monolayer remodeling is driven by local enhanced traction force after apoptosis, which triggers the process of fluidization and mesenchymal-like migration. Our results show the quick recovery of epithelial homeostasis when being interrupted by cell apoptosis, and indicate the importance of apoptosis-induced mechanical force in mediating cell behaviors to maintain the structural integrity of epithelium.

## Introduction

Epithelial monolayers line most surfaces and internal cavities of the body, which act as dynamic barriers between the internal and the external environment. Maintaining the integrity and homeostasis of epithelium is of central importance to an organism’s survival^1^. However, the homeostasis of epithelium can be disrupted under many circumstances. For example, a wound, usually generated by sudden mechanical damages, will result in epithelial integrity disruption. Numerous studies investigating the process of wound healing in epithelial monolayers have revealed that when the protective barrier is broken, a regulated sequence of biochemical and biophysical events is initiated to repair the sudden damage^2,3^. Another natural interruption to the homeostasis of epithelia is cell apoptosis, which exists ubiquitously in multicellular organisms. Unlike the wound, apoptosis is a highly gene-regulated process used by multicellular organisms to dispose unwanted cells, and the self-destruction of a cell is typically characterized by a series of morphological features, such as cell shrinkage and fragmentation into membrane-bound apoptotic bodies. Thus, the cellular events of apoptosis might be expected to challenge epithelial integrity.

Nonetheless, epithelial barriers can remain intact when apoptosis is stimulated^4^. It has been shown that the integrity of epithelial monolayers can even be preserved after triggering apoptosis of more than 50% cells by UV^4^. Thus, mechanisms must exist to preserve epithelial integrity in the face of the inevitable impact of apoptosis. For instance, under physiological conditions, intestinal epithelium has a very high rate of cell apoptosis^5^; correspondingly, a very high rate of cell renewal is observed to maintain the equilibrium of cell numbers^5^. However, maintenance by cell renewal would have low efficiency especially when cells divide slowly or when a massive disruption of epithelium occurs.

In this work, we establish an inducible apoptosis system and investigate the response of epithelial monolayer to induced apoptosis. We show that the homeostasis of structural integrity in epithelial monolayer is severely interrupted by induced cell apoptosis with increasing defect area and permeability to small molecules across the epithelium. To achieve a quick restoration of the structural and functional integrity, cell spreading area, instead of cell number, is significantly increased within a few hours after induction of apoptosis. Meanwhile, cell migratory speed suddenly increases, which allows a rapid and global reorganization of the monolayer. These morphological and migratory changes are attributed to the increased traction force induced by the apoptotic cells within the monolayer. Inhibition of the forces using blebbistatin prevents the spreading of the cells and the acceleration phase of cell migration as well as the restoration of structural and functional integrity. Taken together, our results reveal that the integrity of epithelial monolayers is quickly recovered by cell spreading and cell redistribution after apoptosis, and underline the importance of mechanical force-driven migration as a quick-response mechanism in the apoptosis-disrupted epithelial monolayers.

## Results

### A cell system is constructed to induce apoptosis

MCF10A cell line, derived from human non-tumorigenic breast epithelium, has been widely used as a model system to study the barrier function of epithelial tissues^6–8^. We genetically modified the wild-type MCF10A cell line (wt-MCF10A) with a chemical responsive caspase9 system, so that the apoptosis pathway can be activated by a small-molecule inducer and results in cell apoptosis^9^ (Fig. 1A). The inducible MCF10A cell line is referred as ind-MCF10A here. To validate whether the inducer can efficiently trigger the apoptosis of ind-MCF10A cells, we add 25 nM inducer to both wt-MCF10A and ind-MCF10A cells. While no apoptosis is observed in the wt-MCF10A cell population (Fig. 1B, left panel), almost all the ind-MCF10A cells undergo apoptosis within 100 min (Fig. 1B, right panel). To better monitor cell behaviors, both wt-MCF10A and ind-MCF10A cells are transfected with a nuclear localization signal tagged green fluorescent protein (NLS-GFP)^10^, making nucleus trackable by green fluorescence. The fluorescence maintains in cell nucleus but disappears when cells undergo apoptosis (Fig. 1B, right panel). Thus, we can quantify the number of survival cells by counting the cell nucleus with bright green fluorescence. Consistent with observations in bright-field images, after adding apoptosis inducer, the number of wt-MCF10A cells does not change, while the percentage of ind-MCF10A cells dramatically reduces to ∼20% within 100 min (Fig. 1C).

**Figure 1.**
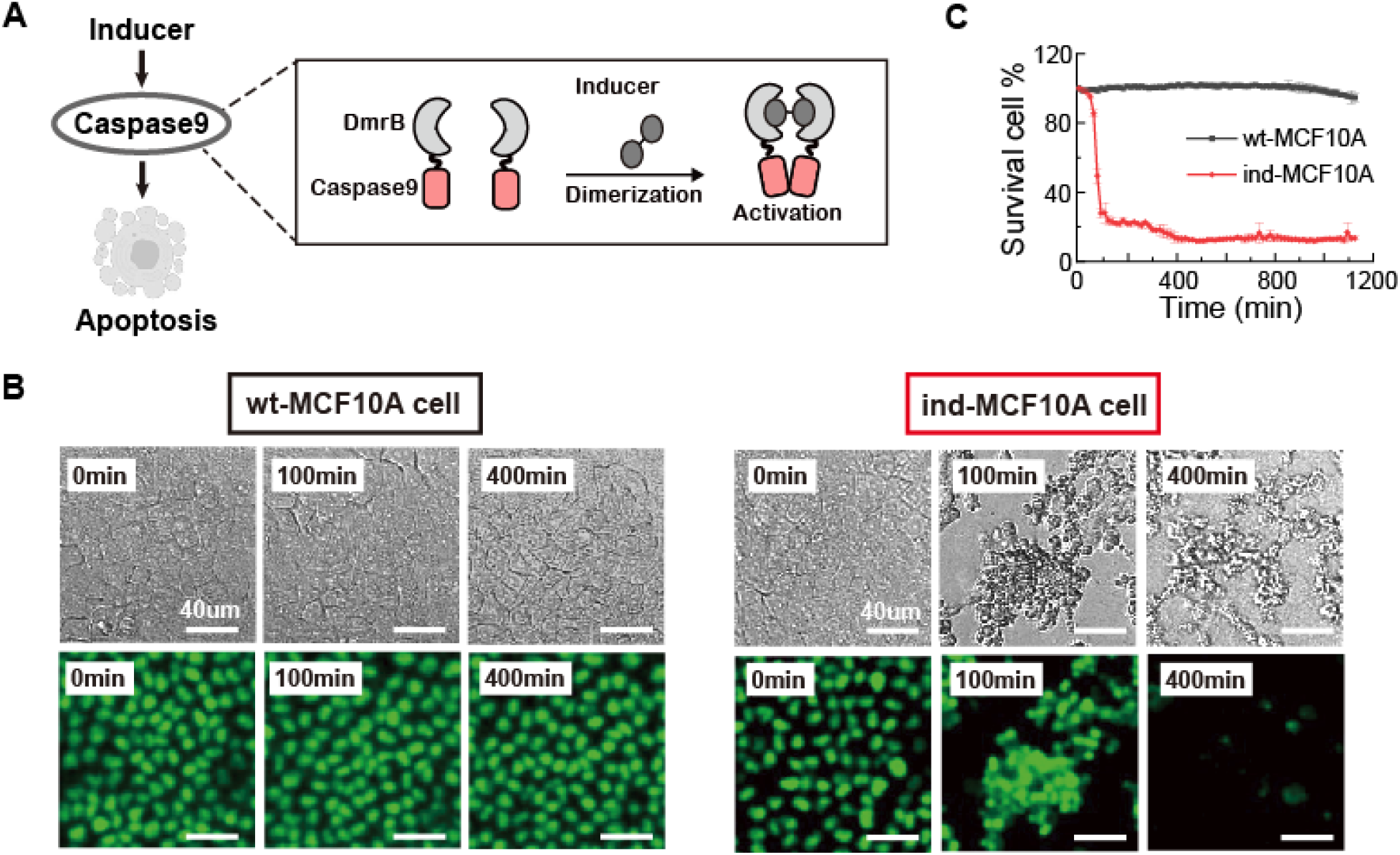
Construction of an inducible system for apoptosis. (**A**) Schematics illustrate the mechanism of inducible apoptosis. (**B**) Microscopy images of BF (upper panel) and NLS-GFP channel (lower panel) for wt-MCF10A cells (left panel) and ind-MCF10A cells (right panel) after being treated with 25 nM inducer for 0 min, 100 min, and 400 min. Scale bar = 40 *µ* m. (**C**) Percentage of survival cells in wt-MCF10A cell population (shown in black line) and ind-MCF10A cell population (shown in red line) after being treated with 25 nM inducer.

### Enhanced cell spreading rather than cell proliferation dominates the quick healing process

To investigate the responses of epithelial monolayer to apoptosis, we first grow an intact epithelial monolayer on a petri dish by mixing ind-MCF10A cells and wt-MCF10A cells at a ratio of 1 to 2. We label ind-MCF10A cells with CellTracker Deep Red dye to distinguish them from the wt-MCF10A cells (Fig. 2A, T=0 min). Then 25 nM inducer is applied to trigger cell apoptosis. As expected, all the ind-MCF10A cells undergo apoptosis, and structural defects are developed near the apoptotic cells (Fig. 2A and Movie S1). Since the structural integrity of the epithelial monolayer is closely related to barrier function, we measure the permeability of the epithelial monolayer and investigate its changes in response to cell apoptosis. We seed four intact monolayers on the microporous membranes of four transwell inserts, and treat them with 25 nM apoptosis inducer for 0 min, 100 min, 1200 min and 2400 min, respectively (Fig. 2B). Then FITC-dextran (4 kDa) is added on the top compartment of each monolayer. To characterize the barrier function, we monitor the FITC-dextran fluorescence intensity in the bottom compartment for one hour using confocal microscope; its changes can be used to quantify the diffusion rate of FITC-dextran across the monolayer. We normalize the diffusion rate by the measurement of the monolayer treated with apoptosis inducer for 0 min. While the normalized diffusion rate of FITC-dextran remains low for the intact epithelial monolayer, cell apoptosis significantly increases the diffusion rate at 100 min and 1200 min (Fig. 2C). However, the diffusion rate at 2400 min decreases to a level similar to that of 0 min and the control, indicating a fully recovery of barrier function of the epithelial monolayer.

**Figure 2.**
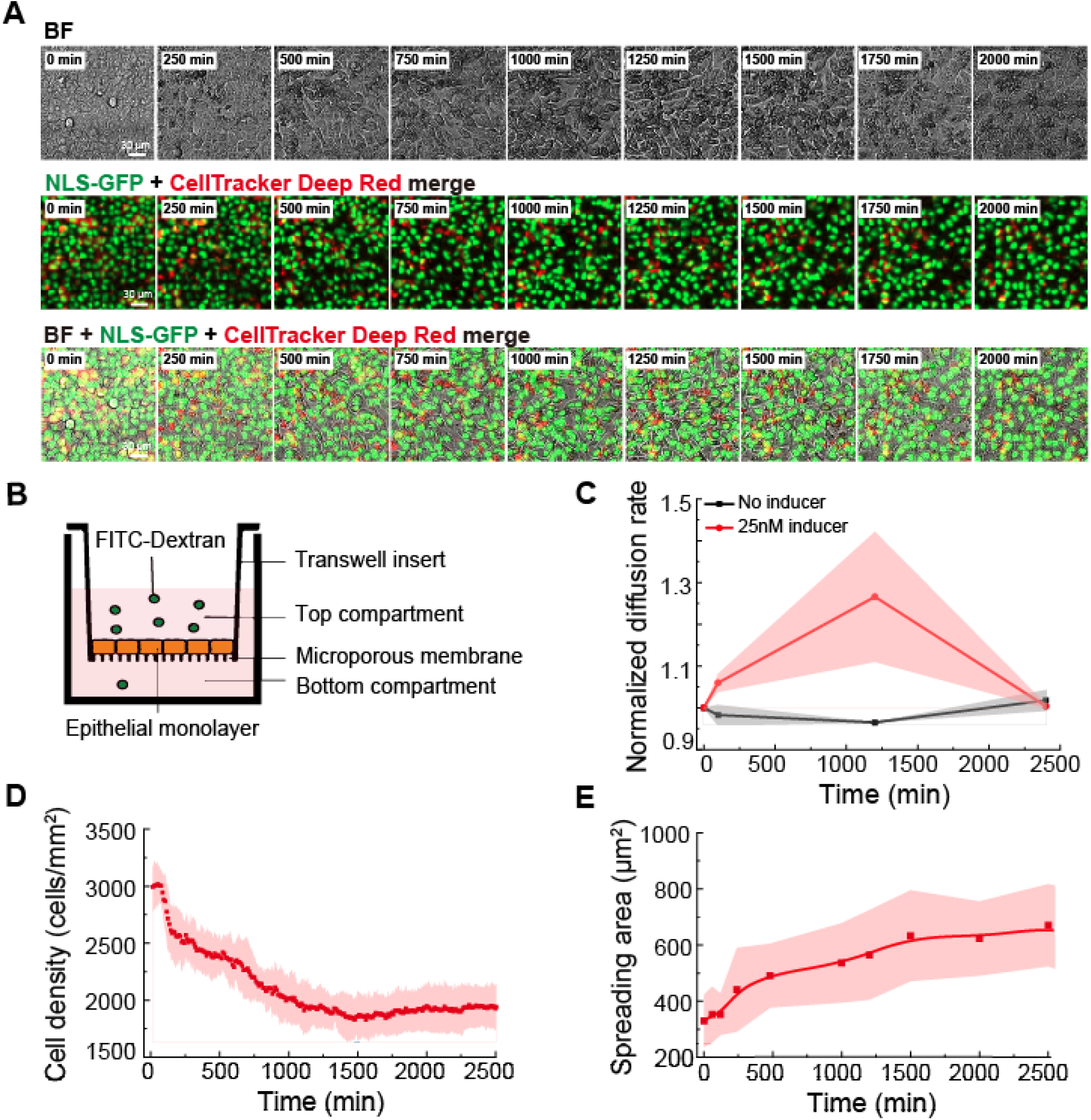
Quick reconstruction of the epithelial monolayer integrity. (**A**) Microscopy images of BF (upper panel), merged NLS-GFP and ind-MCF10A channel (middle panel), and merged BF and NLS-GFP and ind-MCF10A channel (lower panel) for epithelial monolayer after being treated with 25 nM inducer. Scale bar = 30 *µ* m. (**B**) Schematic view of the permeability experiment. Epithelial monolayer is seeded on a porous membrane. The 4kDa FITC-Dextran is added to the top compartment, and the intensity of FITC-Dextran that passes through to the bottom compartment is measured using confocal microscopy. (**C**) Normalized diffusion rate of FITC-dextran across the epithelial monolayers treated without (black line) or with 25 nM apoptosis inducer (red line) for 0 min, 100 min, 1200 min and 2400 min. (**D**) The density of survival cells in the epithelial monolayer after being treated with 25 nM inducer. (**E**) The spreading area of survival cells in the epithelial monolayer after being treated with 25 nM inducer.

It has been shown that maintaining the barrier function of epithelium requires a balance between cell death and cell division^11^, and the apoptotic cells may secrete mitogenic signals to induce proliferation of their neighboring cells^12–15^. However, we do not observe a significant increase of cell number within 2400 min after induction of apoptosis (Fig. 2D), at which time window the recovery of barrier function happens; this indicates that the quick healing process is not resulted from cell proliferation. Though the cell number does not increase during the quick healing process, it gradually rises to the initial level at day 5 after triggering apoptosis (Fig. S1), which implies the importance of cell proliferation in maintaining functions of epithelia in the long term. In short term, however, if the cell number does not increase, then each cell has to increase its size to cover the apoptosis-induced defects to maintain the barrier function of the monolayer. Thus, we quantify cell spreading area from time-lapse imaging of the epithelial monolayer after adding apoptosis inducer. Indeed, the average cell spreading area dramatically increases from ∼300 m^2^ to ∼600 m^2^ within 2400 min after the induction of apoptosis (Fig. 2E). As cells in the initial intact monolayer are closely packed together, the increased cell spreading area is likely due to the space released by apoptotic cells.

Taken together, using a functional epithelium formed by mixing ind-MCF10A cells with wt-MCF10A cells, we find that the integrity of the epithelial monolayer is interrupted by the induction of cell apoptosis. However, the interrupted monolayer can quickly heal through increasing cell spreading area rather than cell renewal.

### Mesenchymal-like cell migration enables the reorganization of the monolayer structure

We next investigate the dynamics of the healing process. Time-lapse imaging of the epithelial monolayer after being treated with apoptosis inducer shows that the migration pattern of cells changes over time (Movie S1). To quantify this effect, we track positions and trajectories of each single cell (Fig 3A), and analyze their migration speed (Fig. 3B, averaged by 20 cells). A significantly enhanced migratory dynamics is observed after induction of apoptosis. The migration speed peaks at ∼1200 min after apoptosis induction, but becomes sedentary at 2400 min. To characterize the migratory dynamics, we calculate the mean square displacement (MSD) of cells with regard to the lag time. The exponent α indicates a sub-diffusive random motion of cells in the very early stage (Fig. 3C, T <= 200 min) and at the end of the quick recovery (Fig. 3C, T >= 2000 min), but a super-diffusive behavior in the middle stage (Fig. 3C, 200 min < T < 2000 min). Concomitantly with the changes of migratory dynamics from a solid-like to a fluid-like phase, we also observe an obvious shift of cell shape from round to elongated at ∼1200 min after adding apoptosis inducer, and the shape eventually turns back to round after 2000 min (Fig. 3D). This cell shape change can be further quantified by calculating the shape index (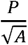, A, area; P, Perimeter) shown in Fig. 3E.

**Figure 3.**
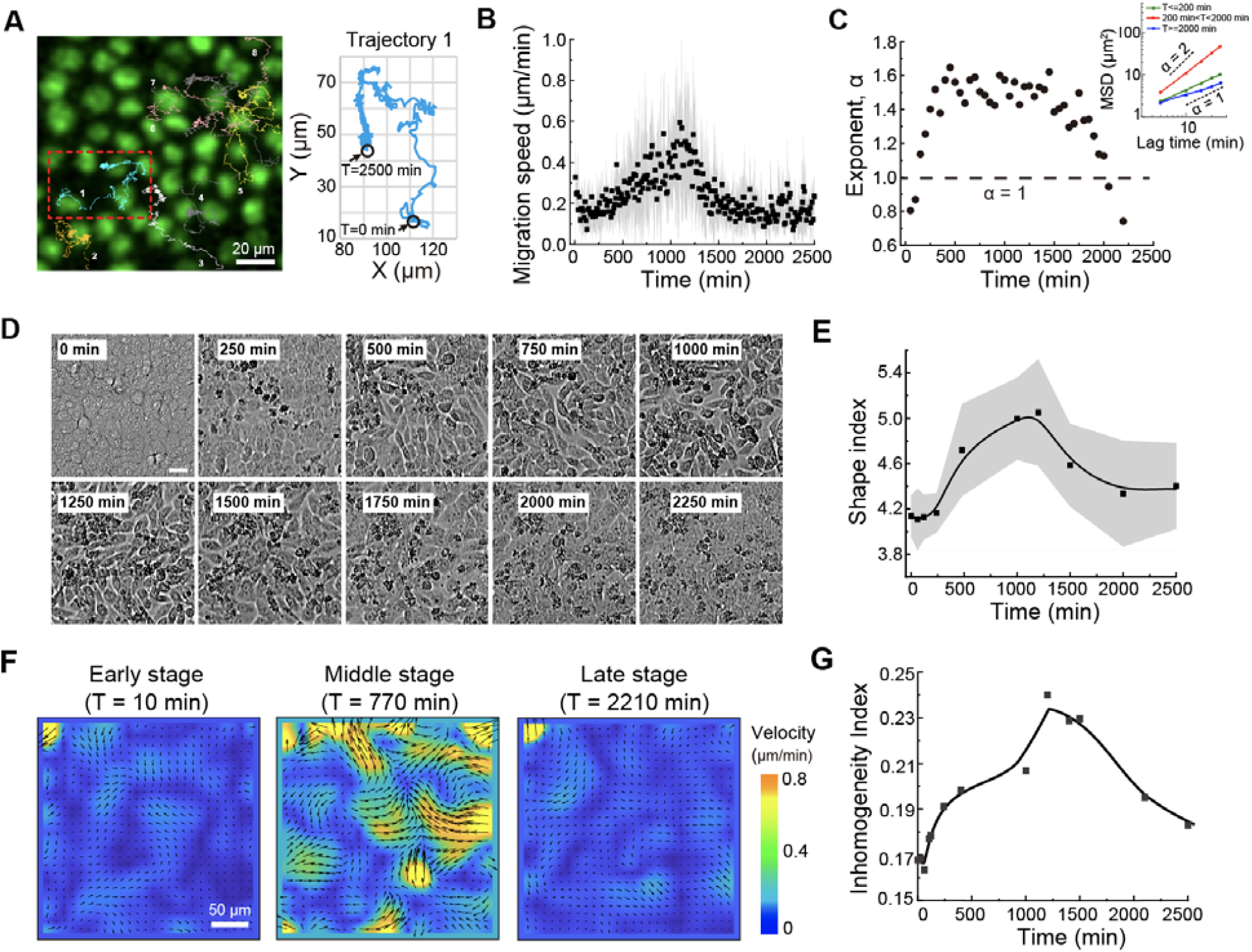
Enhanced migratory dynamics of the epithelial monolayer after cell apoptosis. (**A**) Representative cell trajectories in the epithelial monolayer after being treated with 25 nM inducer. Scale bar = 20 m. (**B**) Cell migration speed in the epithelial monolayer after being treated with 25 nM inducer (averaged by 20 cells, data are shown as mean □±□ standard deviation). (**C**) Exponent of MSD over time. Inner image showing MSD in the early (T <= 200 min, green line), middle (200 min < T <2000 min, red line) and late (T >= 2000 min, blue line) stage. (**D**) Microscopy images of bright field for epithelial monolayer after being treated with 25 nM inducer. Scale bar = 30 μm. (**E**) Shape index of cell bodies in the epithelial monolayer after being treated with 25 nM inducer. (**F**) Heat map of PIV analysis showing epithelium migration in the early (T <= 200 min), middle (200 min < T <2000 min) and late (T >= 2000 min) stage of the recovery process. Black arrows indicate migration direction, and the length of arrows indicates migration velocity. (**G**) Inhomogeneity index is plotted against time after adding 25 nM apoptosis inducer.

The globally changing cell behaviors including cell shape and migratory dynamics is reminiscent of fluidization process^16^, which has been reported as the case in tissue morphogenesis^17^. In our system, PIV (particle image velocimetry) analysis of the epithelial monolayer shows massive flows in the middle stage (200 min < T < 2000 min) after induction of apoptosis (Fig. 3F). Since apoptosis generates defects in the monolayer, the distribution of cells becomes inhomogeneous: cells near the apoptosis spots are sparse, while in the other sites are dense. As we observe, the flow is initiated from the site where apoptosis occurs (Movie. S1), and expands to the whole epithelium, allowing a rapid cell redistribution of homogeneity in the following stage. To quantify this redistribution process, we analyze the inhomogeneity of cell distribution in the monolayer. We define an “inhomogeneity index” as the standard deviation of the nearest neighbor distance (NND) of each single cell in the monolayer divided the mean NND. Indeed, the inhomogeneity index has a sharp increase within 200 min when most apoptosis events happen, while it drops to the similar level as the initial status after massive flows happen (Fig. 3G), suggesting a cell redistribution during the quick healing process triggered by cell apoptosis.

Taken together, enhanced migratory dynamics of epithelial monolayer which initiated from the apoptosis sites, enables the reorganization of the monolayer structure and a rapid recovery of epithelial integrity.

### Increased cell migration is driven by apoptotic force

It has been reported that the forces exerted by apoptotic cells (apoptotic force) drive the tissue dynamics during embryonic development^18,19^, therefore we investigate the effects of cell apoptosis on the dynamics of traction forces generated by the epithelial monolayer. We seed the epithelial monolayer on a soft polyacrylamide (PA) hydrogel (shear modulus: 150 pa) embedded with fluorescent tracer particles, thus the deformation field induced by apoptotic forces can be measured by tracking the motion of these tracers before and after cell apoptosis. The deformation field is relatively homogeneous before apoptosis (Fig. 4A, T=0 min), and a sudden increase in local deformation is observed at random spots at 100 min after the induction of apoptosis (Fig. 4A, T=100 min). These deformations then transit into a global mode (Fig. 4A, T=1200 min) and eventually stabilize at a relatively homogeneous status (Fig. 4A, T=2400 min). In addition to the spatial evolution of the deformation field (Fig. 4C, lower panel), the average deformation presents a similar trend to the average migration speed, with a remarkable increase after triggering apoptosis for 1200 min and finally stabilizes at the level similar to the initial stage (Fig. 4B and Fig. 3B). These results suggest that an increase in traction forces is associated with the global migratory phase in our system. Moreover, the transition from a quiescent and solid-like phase to a hypermobile and fluid-like phase also suggests a change in cell-cell mechanical interaction. Indeed, by calculating the spatial correlation function of the deformation field^20^ at different stages (Fig. S2A), we find that the correlation length is also surged in the middle stage (200 min < T < 2000 min) (Fig. S2B), supporting a long-ranged cell-cell interaction. The increase in correlation length agrees well with the collective and global migratory dynamics (Fig. 3B).

**Figure 4.**
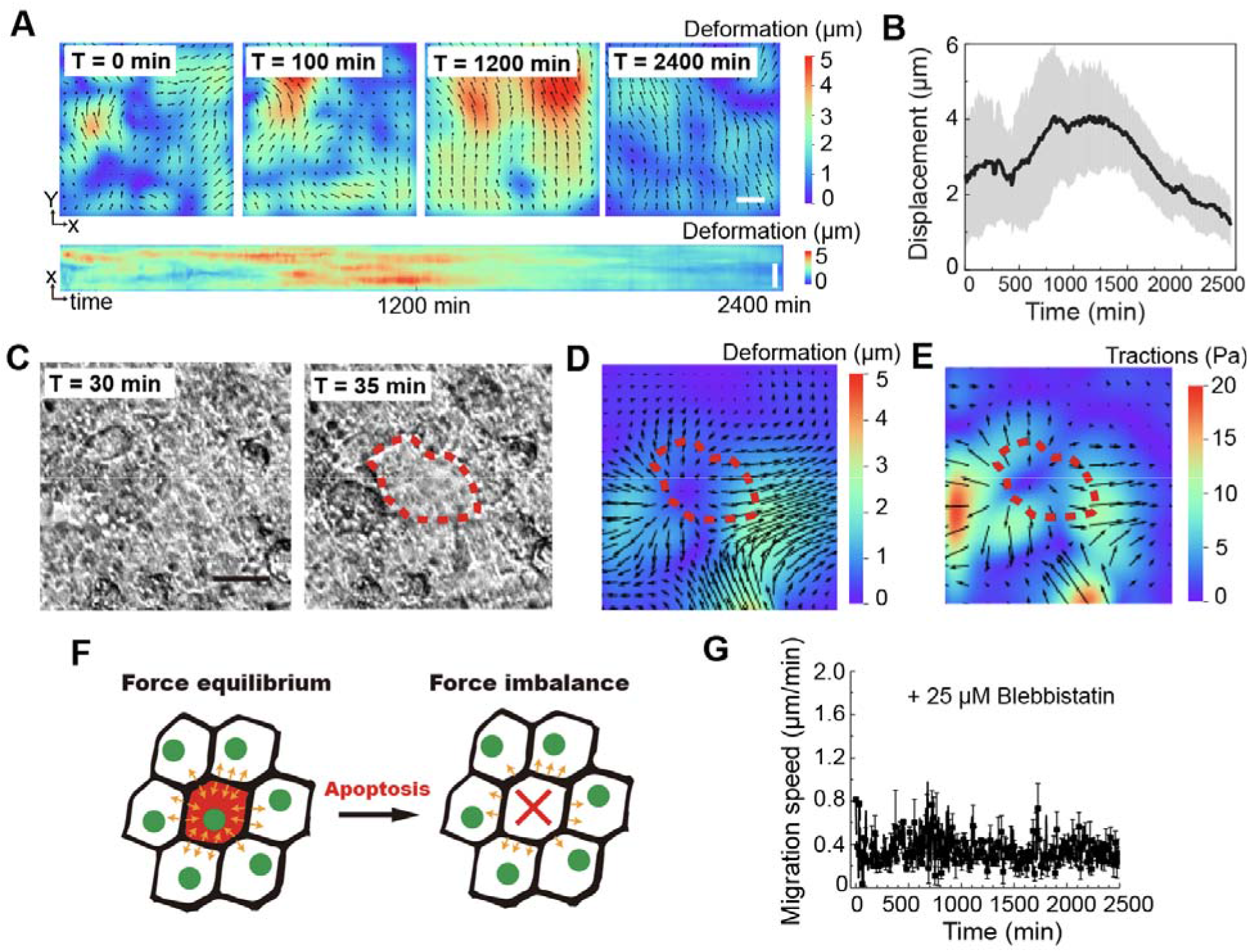
Apoptotic forces drive cell migration in epithelial monolayer after apoptosis induction. (**A**) Deformation field induced by apoptotic forces after adding apoptosis inducer for 0 min, 100 min, 1200 min and 2400 min (Upper panel: scale bar = 50 m); The Kymograph showing changes of substrate deformation over 2400 min after adding 25 nM inducer (Lower panel: scale bar = 20 μm). (**B**) Displacement of matrix over time after adding 25 nM inducer. (**C**) Bright field image showing the development of the apoptosis-induced wound in the epithelial monolayer. Red line indicates the position of the wound. Scale bar = 25 m. (**D**) Local deformation field around the apoptosis-generated wound area. (**E**) Traction force around the apoptosis-generated wound area. (**F**) Schematic of contraction force in apoptosis interrupted epithelium. (**G**) Cell migration speed in the epithelial monolayer after being treated with 25 nM inducer and 25 µM blebbistatin (Data are shown as mean□±□standard deviation).

Interestingly, in the beginning (T<=200 min) where most of apoptotic events occur, we observe a small peak in the average deformation (Fig. 4B), which indicates a direct link between the cell apoptosis and local increases in the deformation. To quantify the local changes in the matrix deformation during an apoptotic event, we calculate the relative deformation field before and after cell apoptosis. The apoptosis generates a local wound in the monolayer within 5 min (Fig. 4C-4E, indicated by the red dash circle) as well as a local matrix deformation pointing outward from the apoptotic cells, which is consistent with the cell migration direction. Furthermore, we calculate the absolute changes in traction stress based on the relative matrix deformation with previously established methods^21,22^, and find that the apoptotic event indeed leads to an increase in traction stress (Fig. 4E). These results led us to propose that the induction of apoptosis breaks the force equilibrium of epithelial monolayer, which dictates a balance of tensile forces between neighboring cells (Fig. 4F). Therefore, remaining live cells will get a net increased traction stress and migrate. To validate this hypothesis, we pretreat cells with blebbistatin to inhibit myosin II activity, as tensile force in cells is generated by actomyosin networks through the activity of myosin motors. As a result, the cell migration speed is no longer increasing after induction of apoptosis (Fig. 4G).

## Discussion

Apoptosis exists widely in epithelial tissues and often interrupts the homeostasis of epithelium integrity. However, maintaining the structural integrity is crucial for epithelial monolayers to perform the protective function as a physical barrier. In this study, we investigate the effect of cell apoptosis on epithelial monolayer and the mechanism of how epithelium maintains the structural integrity after apoptosis. We show that an intact epithelial monolayer is disrupted by induced cell apoptosis with increasing permeability to small molecules across the epithelium. Interestingly, the monolayer heals quickly within a few hours, during which time the cell number remains unchanged while cell spreading area significantly increases. As cells in the initial intact epithelial monolayer are closely packed, the increased cell spreading area is presumably due to the released space by apoptotic cells. Moreover, we observe a rapid and global reorganization of the epithelial monolayer through vigorous cell migration after triggering apoptosis; the movement of epithelial monolayer facilitates the recovery process. Therefore, the increased cell spreading together with the enhanced cell migration ensures a rapid reconstruction of the epithelium integrity.

Previous work has shown that enhanced mechanical force can help cells overcome their surrounding energy barrier^23^ and thus allow cells as a whole to migrate and flow. In our system, we propose that the induction of apoptosis breaks the force equilibrium of epithelial monolayer. As a result, the remaining live cells experience a net increased traction stress and thus migrate. We validate the proposal by treating cells with myosin inhibitor blebbistatin^24^, as actomyosin-mediated contractility is a highly conserved mechanism for generating mechanical stress in animal cells^25^. In this case, the epithelial monolayer does not show enhanced migratory dynamics any more upon the induction of apoptosis, indicating that the increased cell migration is driven by actomyosin-mediated traction force.

To conclude, as a quick response to induced apoptosis, the epithelial monolayer maintains its structural integrity by cell spreading and migrating. Cell spreading ensures an adequate area to fill the apoptosis-generated defects, while the traction force-driven migration enables a global reorganization, making it more efficient to keep the monolayer integrity.

## Supporting information

Supplementary informatioin

## Acknowledgement

We acknowledge the support of the National Key Research and Development Program of China (Grant No. 2020YFA0908200) and the National Natural Science Foundation of China (Grant No. 12090054). The work at Harvard was supported in part by the NIH (Award number 60051124 HU).

## Author Contributions

Yuan Yuan: investigation (lead), conceptualization, formal analysis, writing original draft and review & editing. Yu Long Han: investigation (lead), conceptualization, formal analysis, and draft review & editing. Jing Xia: investigation (lead), conceptualization, formal analysis, and draft review & editing. Hui Li: investigation (supporting) and formal analysis. Chao Tang: resources and draft review & editing. Fangfu Ye: funding acquisition, supervision, and draft review & editing. David A. Weitz: conceptualization, funding acquisition, supervision, and draft review & editing. Ming Guo: conceptualization, supervision, and draft review & editing.

## Materials and Methods

### Cell culture

MCF10A cells are cultured in a DMEM/F12 medium (Invitrogen, 11965-118) supplemented with 5% horse serum (Invitrogen, 16050-122), 20 ng/ml epidermal growth factor (Peprotech, AF-100-15), 0.5 μg/ml hydrocortisone (Sigma, H-0888), 100 ng/ml cholera toxin (Sigma, C-8052), 10 μg/ml insulin (Sigma, I-1882) and 1% penicillin and streptomycin (Thermo Fisher, 15140122). The cells are collected using a 0.05% trypsin-EDTA solution (Thermo Fisher, 25300054) after they reach confluence within T-25 flasks in a standard cell culture incubator.

### Apoptosis induction

The ind-MCF10A cell line is constructed based on wt-MCF10A cells. The wt-MCF10A cells are transfected with DmrB-Caspase9 using lentivirus, which is pre-generated from 293T cells: Three lentivirus vectors pHR-mCherry, pCMVdR8.91, and pMD2.G are co-transfected into 293T cells for lentivirus package and generation. The ind-MCF10A cells with DmrB-Caspase9 are selected in a puromycin-containing culture medium (0.4 mg/ml; Thermo Fisher, A1113802). We use 25nM apoptosis inducer AP20187 (Takara, 635059) to trigger the apoptosis of ind-MCF10A cells.

### Time-lapse imaging of cell migration

To visualize the cell nuclei, both wt-MCF10A and ind-MCF10A cells are transfected with a nuclear localization sequence tagged green fluorescent protein (NLS-GFP)^10^, making nucleus trackable by green fluorescence. Cells were plated in 24-well plate (In Vitro Scientific) with a density of 80,000 per well and were imaged in an incubation chamber for 48 h with a time interval of 5 min. All images were captured by a confocal microscope (Leica, TCL SP8) equipped with 10×/1.3 objective lens. We used filter sets that are optimized for the detection of GFP and Deep Red fluorescence. GFP fluorescence was excited at 488 nm and collected at the wavelength range from 500 nm to 550 nm. Deep Red fluorescence signal was excited by 633 nm laser and collected at the wavelength range from 640 nm to 680 nm.

### Cell migration analysis

The movement of each individual cell is analyzed by tracking the nucleus in green fluorescent channel with particle tracker plugin of ImageJ (https://imagej.nih.gov/ij/). The extracted trajectories are then analyzed with a customized code in MATLAB (Mathworks, MA, USA) to calculate the velocity and mean-square-displacement (MSD) of cell migration.

### Cell number and cell morphology analysis

Cell number was counted with ‘Analyze particle’ function of ImageJ based on the GFP fluorescence signals of cell nucleus. For cell morphological measurements, cell shape and cell spreading area was measured based on bright field images. Circularity (C = 4rrA/P^2^, A, area; P, Perimeter) was used to describe cell shapes. Cell nucleus area was quantified from the fluorescent images of the nucleus.

### Permeability experiment

The ind-MCF10A cells and wt-MCF10A cells are mixed at the ratio of 1:3 and seeded on a microporous membrane of a transwell to form an epithelial monolayer. The 4 kDa FITC-dextran is added on the top compartment. After one hour, we measure the fluorescence intensity of the bottom compartment to determine how much the FITC-dextran permeates through. We use an epithelial monolayer without being treated with apoptosis inducer as the negative control and compare it with that of the monolayer being treated with apoptosis inducer for 0 min, 100 min, 1200 min and 2400 min, respectively.

### Traction force microscopy

A solution for surface activation is made of 1 mL APTES, 1 ml 1M NaOH and 98 mL 100% Ethanol. The glass-bottom 24-well plate is activated by soaking in the solution at room temperature for 10 min, followed by three times wash with 100% ethanol and ddH2O, respectively. Then PA gels with shear modulus of 150 Pa were made on the air-dried 24-well plate: For each 2.5 mL PA gel solution, 1 μl of 1 μm fluorescent beads (Sigma, L9654) were added along with 15 μl 10% APS and TEMED and 10 μl TEMED to achieve a final bead concentration of 5 × 10^10^/ml. Then 38 μl of the gel mixture was added to one well of the 24-well plate and covered with a 15 mm cover glass. We quickly inverted the plate to ensure sedimentation of beads on the surface of PA gels. After solidification, the gels were then treated with sulfo-SANPAH and coated with collagen type-I (Advanced BioMatrix, #5005) for overnight. Then the cells were seeded on the collagen-coated PA gel in 24-well plate and images were taken under the confocal microscope mentioned above. Fluorescent beads was excited at 520 nm and collected at the wavelength range from 530 nm to 550 nm. pyTFM was used to calculate the bead displacement and traction force^21,26^.

## References

1. Macara, I. G., Guyer, R., Richardson, G., Huo, Y. & Ahmed, S. M. Epithelial homeostasis. Current Biology 24, R815–R825 (2014).

2. Belacortu, Y. & Paricio, N. Drosophila as a model of wound healing and tissue regeneration in vertebrates. Developmental Dynamics 240, 2379–2404 (2011).

3. Ajeti, V. et al. Wound healing coordinates actin architectures to regulate mechanical work. Nature Physics 15, 696–705 (2019).

4. Rosenblatt, J., Raff, M. C. & Cramer, L. P. An epithelial cell destined for apoptosis signals its neighbors to extrude it by an actin- and myosin-dependent mechanism. Current Biology 11, 1847–1857 (2001).

5. Patankar, J. V. & Becker, C. Cell death in the gut epithelium and implications for chronic inflammation. Nature Reviews Gastroenterology and Hepatology 17, 543–556 (2020).

6. Marshall, A. M., Pai, V. P., Sartor, M. A. & Horseman, N. D. In vitro multipotent differentiation and barrier function of a human mammary epithelium. Cell and Tissue Research 335, 383–395 (2009).

7. Stull, M. A. et al. Mammary gland homeostasis employs serotonergic regulation of epithelial tight junctions. Proceedings of the National Academy of Sciences of the United States of America 104, 16708–16713 (2007).

8. Lourenço, A. R. et al. C/EBPL is crucial determinant of epithelial maintenance by preventing epithelial-to-mesenchymal transition. Nature Communications 11, (2020).

9. Yuan, Y. et al. Cell-to-cell variability in inducible Caspase9-mediated cell death. Cell Death & Disease 13, 34 (2022).

10. Han, Y. L. et al. Cell swelling, softening and invasion in a three-dimensional breast cancer model. Nature Physics 16, 101–108 (2020).

11. Young, H. K. & Raphael, Y. Cell division and maintenance of epithelial integrity in the deafened auditory epithelium. Cell Cycle 6, 612–619 (2007).

12. Kawamoto, Y., Nakajima, Y. I. & Kuranaga, E. Apoptosis in cellular society: Communication between apoptotic cells and their neighbors. International Journal of Molecular Sciences 17, (2016).

13. Fan, Y. & Bergmann, A. Apoptosis-induced compensatory proliferation. The Cell is dead. Long live the Cell! Trends in Cell Biology 18, 467–473 (2008).

14. Ryoo, H. D. & Bergmann, A. The role of apoptosis-induced proliferation for regeneration and cancer. Cold Spring Harbor Perspectives in Biology 4, (2012).

15. Brock, C. K. et al. Stem cell proliferation is induced by apoptotic bodies from dying cells during epithelial tissue maintenance. Nature Communications 10, (2019).

16. Park, J. A. et al. Unjamming and cell shape in the asthmatic airway epithelium. Nature Materials 14, 1040–1048 (2015).

17. Petridou, N. I., Grigolon, S., Salbreux, G., Hannezo, E. & Heisenberg, C. P. Fluidization-mediated tissue spreading by mitotic cell rounding and non-canonical Wnt signalling. Nature Cell Biology 21, 169–178 (2019).

18. Monier, B. et al. Apico-basal forces exerted by apoptotic cells drive epithelium folding. Nature 518, 245–248 (2015).

19. Toyama, Y., Peralta, X. G., Wells, A. R., Kiehart, D. P. & Edwards, G. S. Apoptotic force and tissue dynamics during Drosophila embryogenesis. Science 321, 1683–1686 (2008).

20. Angelini, T. E., Hannezo, E., Trepat, X., Fredberg, J. J. & Weitz, D. A. Cell migration driven by cooperative substrate deformation patterns. Physical Review Letters 104, (2010).

21. Butler, J. P., Toli-Nørrelykke, I. M., Fabry, B. & Fredberg, J. J. Traction fields, moments, and strain energy that cells exert on their surroundings. American Journal of Physiology - Cell Physiology 282, (2002).

22. Tambe, D. T. et al. Collective cell guidance by cooperative intercellular forces. Nature Materials 10, 469–475 (2011).

23. Park, J. A., Atia, L., Mitchel, J. A., Fredberg, J. J. & Butler, J. P. Collective migration and cell jamming in asthma, cancer and development. Journal of Cell Science 129, 3375–3383 (2016).

24. Kovács, M., Tóth, J., Hetényi, C., Málnási-Csizmadia, A. & Seller, J. R. Mechanism of blebbistatin inhibition of myosin II. Journal of Biological Chemistry 279, 35557–35563 (2004).

25. Murrell, M., Oakes, P. W., Lenz, M. & Gardel, M. L. Forcing cells into shape: The mechanics of actomyosin contractility. Nature Reviews Molecular Cell Biology 16, 486–498 (2015).

26. Bauer, A., Prechová, M., Gregor, M. & Fabry, B. pyTFM: A tool for Traction Force and Monolayer Stress Microscopy. bioRxiv (2020).

