## Supplementary informatioin for "Recovery of structural integrity of epithelial monolayer in response to massive apoptosis-induced defects"

**This PDF file includes:**

Supplementary Figs. 1 to 2

**Other Supplementary Materials for this manuscript include the following:**

Supplementary Movie 1

**Supplementary Figures**

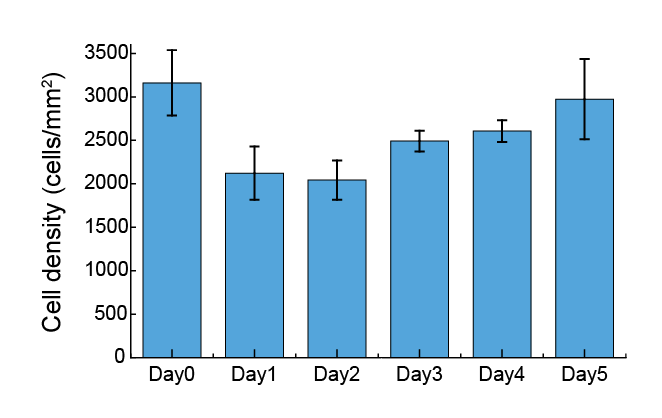

**Figure S1.** Cell density in the epithelial monolayer (ind-MCF10A : wt-MCF10A = 1:2) after addition of apoptosis inducer for different days. Data are shown as mean ± standard deviation.

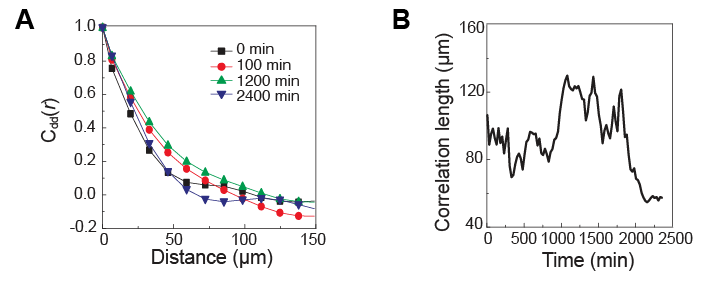

**Figure S2.** (**A**) Correlation function of the deformation field is plotted against distances at 0 min, 100 min, 1200 min and 2400 min. (**B**) Correlation length along with time after adding 25 nM inducer.
